# Poly Pipeline: A Polyvalent Spatial Transcriptomics Workflow Validated Across Polyploid and Diploid Organisms

**DOI:** 10.64898/2026.08.31.748304

**Authors:** Pedro C Carvalho, Tori Millsteed, Robert J Henry

## Abstract

Spatial transcriptomics (ST) has emerged as a transformative approach for visualizing tissue landscapes, yet it faces significant challenges regarding data standardization, sparsity, and the analysis of complex genomes, particularly polyploid plants. To address these limitations, we introduce Poly Pipeline, a robust and universal bioinformatic workflow designed to streamline analysis across diverse plant and animal genomes. The pipeline integrates a comprehensive converter for proprietary formats, clustering algorithms, and hdWGCNA co-expression networks, which indirectly preserves the expression signatures of low-expressed duplicated genes. Benchmarking across datasets from wheat, rice, *Arabidopsis*, and mouse demonstrated the broad applicability of the pipeline in identifying relevant clusters, showing effectiveness across diverse organisms and data types. By providing a unified and reproducible framework, Poly Pipeline addresses a critical gap in analyzing genomic redundancy, especially that related to polyploidy, and promotes FAIR data principles for the broader scientific community.

## Introduction

Spatial transcriptomics (ST) has been recognized as a transformative approach in biological research, enabling the simultaneous quantification of gene expression and the visualization of the tissue landscape at subcellular resolution [1,2]. The development of new technologies involving ST has scaled quickly and exponentially, enabling the construction of comprehensive atlases for different organisms related to a range of factors. However, this has led to problems inherent in work with ‘Big Data’, particularly regarding data management and standardization, processing efficiency and scalability, and high-dimensional sparse datasets [3–7]. The relatively recent development also reflect problems related to proprietary technology, which results in a lack of standardization of the data format output, generating a barrier to generating a universal pipeline for ST [8].

Another critical bottleneck in current ST analysis is the noise related to the sparse data, mainly inherent from the chip technology used but also resulting from low RNA capture efficiency and ‘dropouts’ from poorly designed data processing analysis [9,10]. Recent advances have also explored adaptive Gaussian smoothing to improve signal-to-noise ratios and recover dropped-out transcripts in high-resolution spatial matrices [11]. Furthermore, numerous bioinformatic tools have been developed to mitigate problems inherent in sparse data, including spatial clustering, highly variable gene identification, and cell-type deconvolution, with most of those technologies being focused on mammalian tissues and diploid organisms [12–14]. The development of ST technologies for plants faces several problems, including structural issues such as the rigid cell walls and high starch content, which often lead to limited or diminished quality of the resulting capture efficiency and data output, and subsequent analysis issues, including complex and usually poorly annotated genomes, hindering the mapping and quantification quality [15,16].

The development of new technologies focused on plants, primarily related to non-model and polyploid organisms, however, is still lacking, resulting in a gap in the analysis of these common commercially important organisms. Analysis focused on polyploid organisms faces the deepest problems, including allelic complexity and duplicated gene functions, which may bias the analysis, reinforcing the need for a robust pipeline capable of normalizing the data and mitigating these problems [17]. To address these challenges, we introduce the Poly Pipeline, a novel bioinformatic workflow designed for the robust analysis of spatial transcriptomics data. While validated on complex polyploid plant genomes, its modular design makes it highly effective for analyzing a wide range of diploid and polyploid organisms. To achieve this, the pipeline integrates a concise analytical roadmap, by initially converting different input formats to standardize fragmented proprietary inputs; proceeds with robust quality control and multi-mode spatial clustering to accurately delineate tissue domains; and culminates with the implementation of high-dimensional Weighted Gene Co-expression Network Analysis (hdWGCNA) [18]. By coupling universal data standardization with sparsity-aware network generation, this workflow directly bridges the current bottlenecks of spatial transcriptomics processing to comprehensive biological interpretation.

### Pipeline

The rapid progress of ST technologies has resulted in the development of multiple bioinformatics tools designed to tackle the unique challenges of spatially resolved data. Toolkits such as *Seurat* [19] and *Giotto*[13] were the first developed, offering end-to-end workflows for quality control, dimensionality reduction, and visualization. Although Seurat was paramount to single-cell RNA-seq (scRNA-seq) analysis, it was often limited to basic clustering that did not fully exploit spatial adjacency, and while *Giotto* introduced a more specialized suite of tools, including spatial network construction and the identification of spatial domains, it has been noted for its substantial resource requirements and complex dependencies [3,13,19,20]. Similarly, *Squidpy* [21] and *Scanpy* [22] supply standards for ST analysis, providing scalable infrastructure for neighborhood clustering enrichment However, they often require integration with other tools for advanced tasks like deconvolution or precise domain segmentation.

Specific pipelines have been developed to handle the unique data structures of high-resolution platforms, such as the Stereo-Seq Analysis Workflow (SAW) [23]. The specific SAW pipeline provides an efficient, standard processing stream for Stereo-seq data, handling image registration and generating output files, but it is primarily an upstream processing tool rather than a flexible downstream analytical suite. Other pipelines have been developed, including *FaST* [24], that offers an light alternative for raw data mapping and segmentation, but lacks extensive statistical modeling capabilities required for complex biological interpretation. Additionally, *BayesSpace* [10] offers resolution enhancement through probabilistic modeling, but is computationally intensive and less effective at capturing non-linear expression relationships [9]. To address the limitations of generalist tools, particularly in identifying coherent tissue regions, our pipeline integrates the best of each technology into a unified and global pipeline, highlighting the use of *Stereopy* [3] for large-scale data handling and the *hdWGCNA* [18] for analyzing complex sparse datasets to produce co-expression networks (Figure 1).

**Figure 1.**
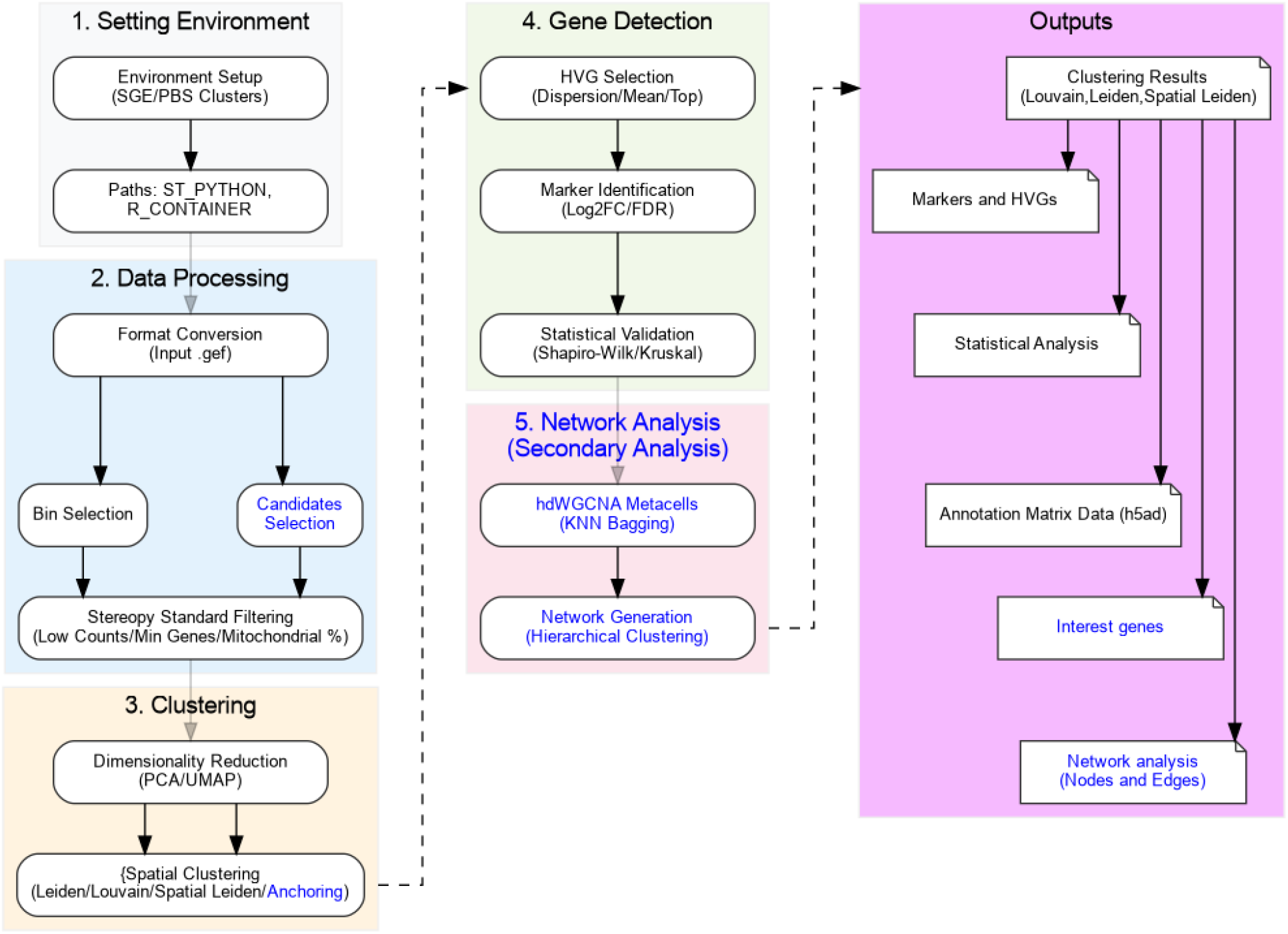
Diagram of the Poly Pipeline workflow, summarizing the sequential steps from environment setup and spatial transcriptomics data processing to clustering, dimensionality reduction, gene detection, and downstream analytical modules. Components and outputs highlighted in blue indicate optional steps that can be activated according to the goals of each analysis.

### Setting environment

The pipeline relies on multiple dependencies from the python and R packages used, however, standard dependencies solutions are not applicable since some of the packages, particularly the *Stereopy* package, are developed for dependencies with specific versions. Furthermore, the analysis requires high computational resources and must be run on high-performance computers (HPCs), which are commonly directed by job scheduler softwares, including Son of Grid Engine (SGE) and Portable Batch System (PBS). Commonly for SGE clusters where the user has no limitations of space or internet access, the environment can be set directly with the provided script from the main pipeline. However, when using PBS clusters, usually the connection to the internet through jobs is limited, and the space provided to users is usually very limited, where the user must install dependencies manually as instructed on the pipeline documentation. Regardless of the method used to install the dependencies, the user must know and provide (variables ‘ST_PYTHON’ and ‘R_CONTAINER’) the correct path for both the python and R environments prior to the execution of the primary analysis, which is provided to the script.

### Data processing

The input data for the pipeline is the bin Gene Expression File (GEF) format, structured with the expression matrices for multiple bin sizes (1, 5, 10, 20, 50, 100, 150, 200) for the complete dataset, i. e., the Molecular Identifier (MID) for each bin of the tissue with the related expression value. The bin size represents the spatial aggregation (resolution) of the sample, by which the DNA nanoball (DNB) patterned arrays, used by the Stereo-seq technology, have a center-to-center distance of approximately 500 to 715 nm [6]. Therefore, while Bin 1 represents the sub-micron physical capture of a single DNB, a bin size of 20 (Bin 20) aggregates adjacent spots to represent a spatial resolution of approximately 10–14 microns, roughly corresponding to the diameter of a single mammalian cell [3]. Complementary segmentation frameworks, such as STCellbin [25], have highlighted the importance of cell boundary and plant cell-wall staining images in defining accurate cellular profiles in Stereo-seq, reinforcing the rationale behind flexible bin aggregation. For this step, the user is able to select the desired bin size (variable ‘BIN_SIZE’) for analysis of data that includes multiple bin sizes, otherwise. The pipeline uses the default bin size of 50, following the documentation instructions, where bin 20 represents one complete mammalian cell, while lower resolutions can introduce noise and higher resolutions result in loss of information [23].

Facing the problem resulting from multiple proprietary technology developed for ST, the pipeline provides a comprehensive converter capable of dealing with most common ST data outputs, including the common annotation matrix files (AnnData - h5ad) and the gene expression matrix (GEM), to the proper GEF format prior to the main analysis. The converter is a secondary analysis where the user provides the file or a complete folder (variable ‘INPUT_PATH’) to be converted to the proper input file prior to the main analysis. Optionally, the user can also define a list of interest genes for differential analysis (variable ‘INTEREST_GENES_PATH’) where they will be analyzed based on an expression threshold (variable ‘EXPRESSION_THR’ or default below 1 Log 2 Fold Change).

The main analysis focuses mainly on the *Stereopy* Package [3] for Python, strictly following the package documentation, filtering low quality bin spots based on the quality control results, filtering bins with low counts (variable ‘MIN_COUNTS’ or default below 20), low number of expressed genes (variable ‘MIN_GENES’ or default below 3) and bins with too many mitochondrial genes (variable ‘PCT_COUNTS_MT’ or default above 5%). The bins are then filtered by region, including only bin spots located within the known limits of the sample (variables ‘MIN_X’, ‘MAX_X’, ‘MIN_Y’, ‘MAX_Y’, or default of the complete tissue).

### Clustering analysis

The clustering analysis of ST data has historically relied on linear models and statistical frameworks to decipher gene expression patterns. Traditional approaches such as Principal Component Analysis (PCA), have been instrumental in dimensionality reduction and the identification of spatial domains [3,26]. More recently, deep learning frameworks utilizing graph attention autoencoders and multiscale deep subspace clustering have emerged to decipher complex spatial domains across diverse platforms [27]. In our analysis, the reduction of dimensionality was performed both with Uniform Manifold Approximation and Projection (UMAP) and Principal Component Analysis (PCA) with the number of principal components established from the elbow plot after the first analysis as previously discussed. Usually, it is recommended to perform a first run with standard variables to generate the custom elbow plot, followed by a second run with the specific value obtained for the generation of the Principal Component analysis (variable ‘N_PCS’ or default 10) (Figure 2). Clusters with high aggregation or with Wilcoxon outlier scores were selected as possible candidates for downstream analysis. The significance of cluster members was determined by the Wilcoxon statistical analysis, a non-parametric test to determine sample bin spots differences similar to the t-test performed for normally distributed samples [28].

**Figure 2.**
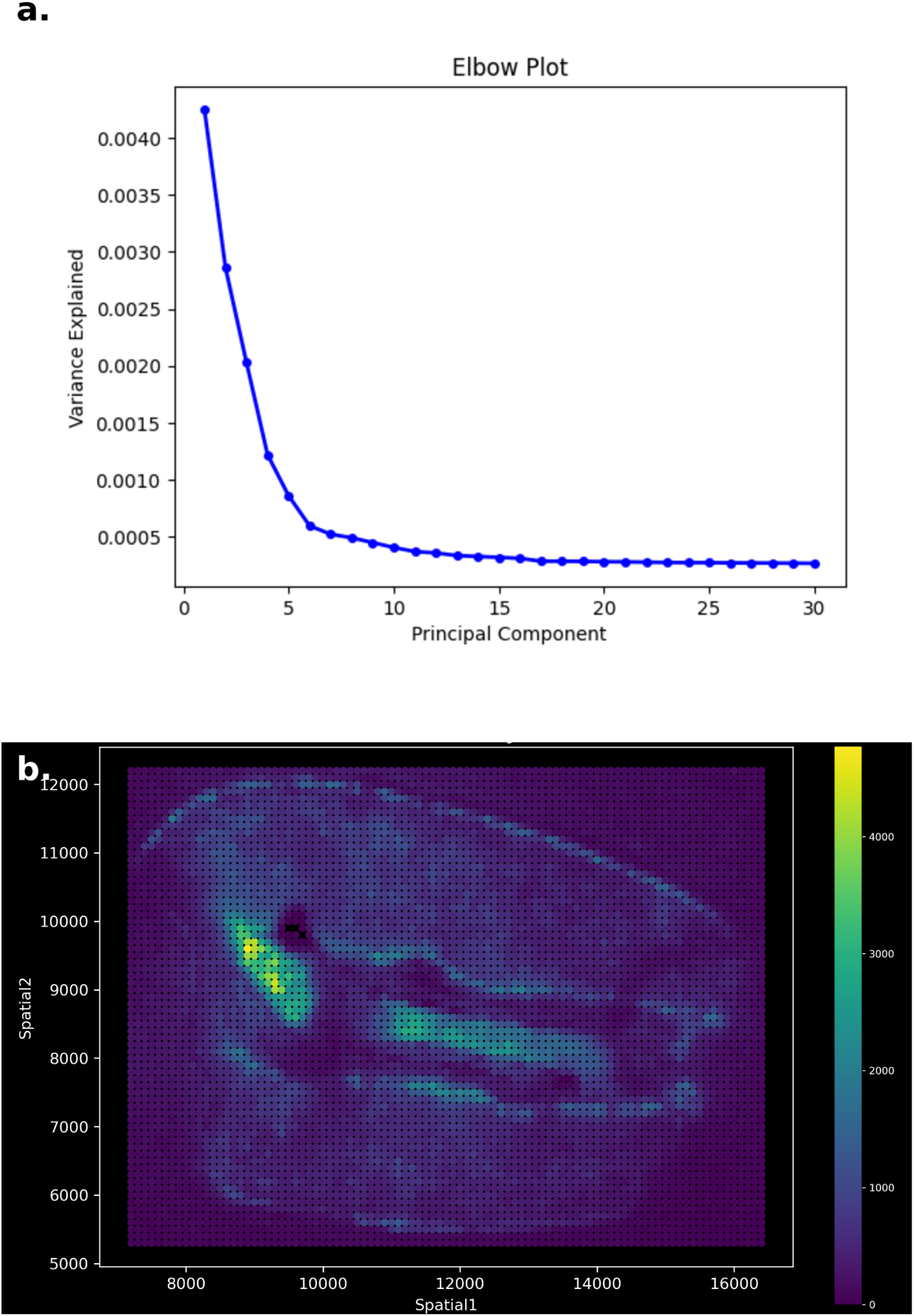
PCA-based dimensionality reduction in the application of the pipeline to wheat spatial transcriptomics data representing (a) Elbow plot showing the variance explained across principal components, used to guide the selection of the number of PCs retained for downstream dimensionality reduction and clustering analyses and (b) Spatial visualization of spot-level expression/count intensity across the wheat tissue section after tissue-only filtering, illustrating heterogeneous molecular signal distribution and regions with increased transcriptional activity. Together, these panels exemplify the PCA-guided analytical module of the pipeline and its integration with spatially resolved expression visualization in wheat data [29].

The clustering is followed by the standard Louvain, Leiden and Spatial Leiden methods, generating distinct results with all associated statistical analyses. This step is improved in the present pipeline with the custom anchoring secondary analysis method, where the user defines marker loci with expression in known regions of interest and all other loci are mapped over them, determining their region by the majority expression over each of the anchors.

### Detection of Highly Variable and Marker Genes

Highly Variable Genes (HVGs) are selected by their dispersion from the mean expression values from the complete samples, being independent of any clustering method applied. The filtering parameters are defined by the user or default based on the suggested documentation, including the minimal mean expression (variable ‘HVG_MIN_MEAN’ or default 0.0125), the maximum mean expression (variable ‘HVG_MAX_MEAN or default of 3) and a minimal dispersion (variable ‘HVG_DISP’ or default 0.5), followed by the selection of the top HVGs (variable ‘HVG_TOP’ or default 2000) for downstream analysis.

For each cluster previously established through the Leiden method all the marker genes were selected based on the *t-test*. The genes were filtered based on the Log 2 Fold Change (including values of 0.5, 1.0 and 2) and by their adjusted *p-values* by the standard Benjamini-Hochberg procedure (False Discovery Rate - FDR) including values lower than 0.05, 0.01 and 0.001 for comparison and validation of the results. All the analyses were supplemented using the *SciPy* package [30] for Python providing statistical significance through the Shapiro-Wilk and Kruskal-Wallis H-tests for both the list of HVGs and marker genes for each individual cluster.

### Network analysis

Traditional Weighted Gene Co-Expression Network Analysis (WGCNA) has long been the gold standard for network analysis in bulk transcriptomics, however, its direct application to ST data faces the inherent problem of sparse data, and that is confounded by the zero-inflated nature of the matrices, leading to fragmented modules and low topological overlap [18,31]. Facing the limitations from WGCNA, our pipeline incorporates the metacell construction and sparsity-aware correlation metrics inherent to high-dimensional Weighted Gene Co-expression Network Analysis (hdWGCNA) [18]. The hdWGCNA package enables the generation of robust gene co-expression modules that incorporate low expressed genes, usually taken as ‘dropout’ on standard methods, characteristic of polyploid organisms with inherent variation in gene dosage [4,15,18,32].

The differentiation from the hdWGCNA lies in the normalization of the data prior to standard WGCNA methods, where highly similar bins are collapsed into ‘metacells’ through the K-nearest neighbors (KNN) bagging algorithm, reducing the number of zero-values in the expression matrix, which generates the major problems of the WGCNA method. After the collapse, the network is generated by the standardized WGCNA procedures including the establishment of pairwise gene correlations, calculation of the weighted correlations using a soft-power threshold (β), and the establishment of network interconnectedness through the Topological Overlap Matrix (TOM), identifying gene modules via unsupervised hierarchical clustering. The expression pattern of an entire module is summarized by its Module Eigengene, defined as the first principal component of the module’s gene expression matrix (Figure 3). The generation of the network is a secondary optional analysis of the pipeline, where the user is provided with all nodes and edges for each individual cluster generated and the complete list, which can be further analyzed.

**Figure 3.**
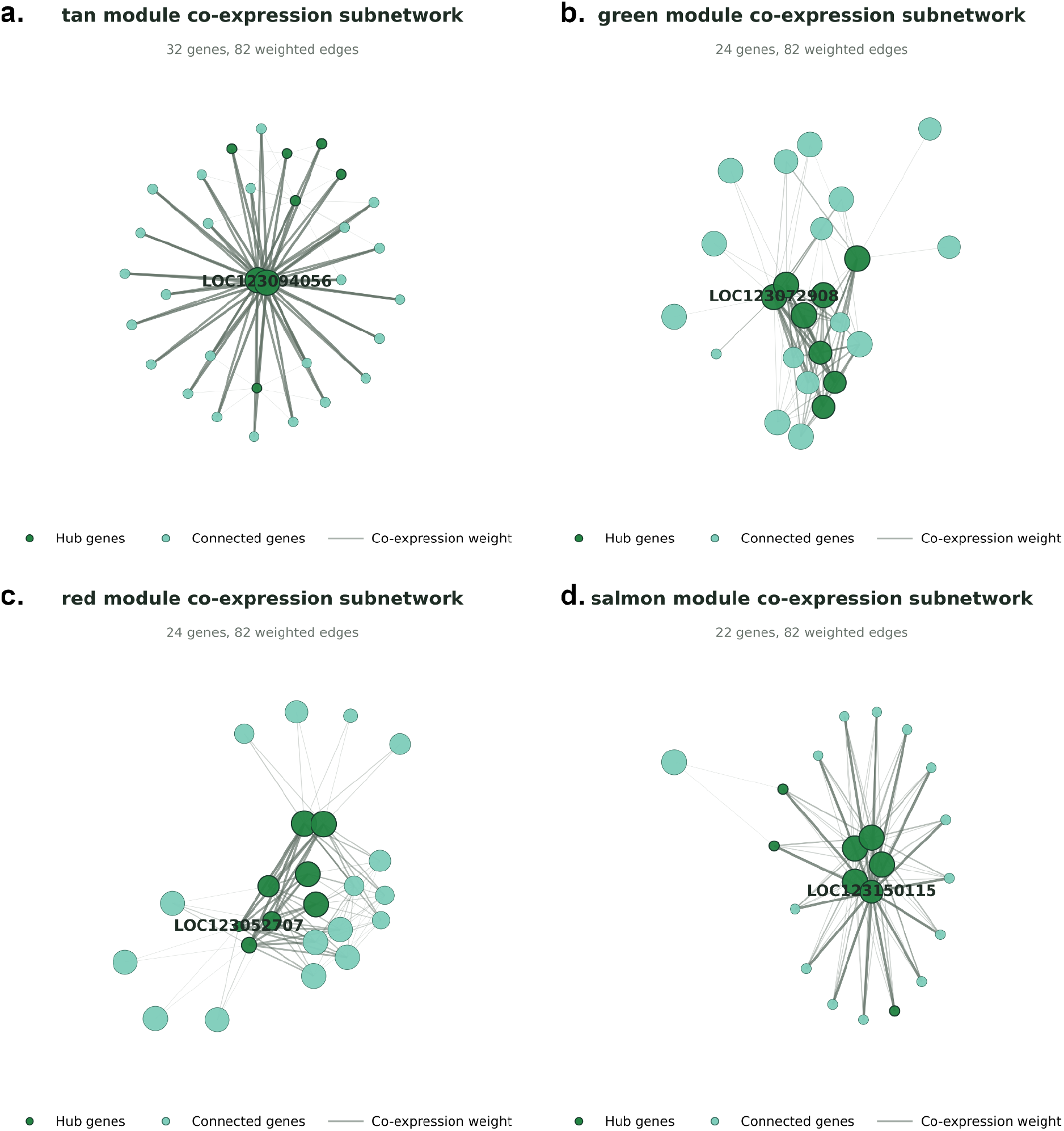
Gene co-expression network construction from the application of the pipeline to wheat spatial transcriptomics data [29]. Representative hdWGCNA-derived module subnetworks generated from node and edge tables exported by the pipeline. a-d, Co-expression subnetworks for the tan, green, red, and salmon modules, respectively. Nodes represent genes, node size reflects module membership strength, darker green nodes indicate hub genes, and edge thickness represents co-expression weight. The visualization illustrates the ability of the pipeline to move from spatial transcriptomics processing to network-level representation of gene modules and candidate hub genes in wheat data.

### Benchmarking

The pipeline was developed based on data obtained from the study of Millsteed et. al. [29] analysis of the developing wheat (*Triticum aestivum*) seed, providing the polyploid data set used for the pipeline development. For this data, the raw analysis was performed with the SAW pipeline [23] which generated the proper input file. However, further benchmarking was performed for multiple organisms, including data for rice [33], *A. thaliana* [34] and mouse [6], where the files were obtained in different formats, proving the efficiency of the converter provided.

The benchmarking results underscore the robustness of the pipeline and its capacity to handle diverse spatial transcriptomics data architectures through its conversion module (Table 1). The ability of the pipeline to successfully process distinct input formats (h5ad and gem) highlights its versatility regardless of the complexity of the input organism. Furthermore, the pipeline demonstrates high versatility by integrating multiple clustering approaches, allowing users to perform standard methods such as Louvain and Leiden alongside spatially-aware algorithms such as Spatial Leiden. Across all tested datasets, the pipeline consistently identified distinct spatial domains regardless of organism complexity or initial data format (Figure 4). While the pipeline successfully executes all clustering modes, the Spatial Leiden algorithm provided a more cohesive spatial division of tissue structures, as evidenced by the 15 well-defined tissue zones identified in wheat compared to the more fragmented 26 clusters generated by the standard non-spatial Leiden method. Even with the high computational requirements of wheat data (40,248 cells and 70,766 genes), the analysis was completed in approximately 23 minutes and 30 seconds, requiring a maximum of 31.8 GB of RAM, confirming that the downstream modules are both computationally efficient and biologically robust.

**Table 1.** Benchmarking Results of the Poly Pipeline Across Different Organisms and Spatial Transcriptomics Data Architectures. Multiple datasets from diverse organisms were selected and tested with the present pipeline and the number of clusters, the time of execution and the RAM consumption were registered for benchmarking purposes.

| Organism | Total<br>Cells | Total<br>Genes | Louvain<br>Clusters | Leiden<br>Clusters | Spatial<br>Leiden<br>Clusters | Time (s) | Peak<br>RAM<br>(GB) |
| --- | --- | --- | --- | --- | --- | --- | --- |
| **Wheat** | 40248 | 70766 | 26 | 26 | 15 | 1411.6 | 31.8 |
| **Mouse** | 4399 | 24238 | 16 | 17 | 13 | 191.2 | 4.4 |
| *Arabidopsis* | 651 | 18266 | 7 | 8 | 5 | 119.8 | 3.2 |
| **Rice** | 1036 | 24301 | 8 | 9 | 7 | 161.7 | 3.5 |

**Table 2.** Statistical results from the Benchmarking Results of the Poly Pipeline Across Different Organisms and Spatial Transcriptomics Data Architectures. The statistical significance of all the analysis were evaluated to provide proper delineation of the pipeline in the identification of highly variable significant genes.

| Organism | Shapiro-Wilk | Skewness | Kurtosis |
| --- | --- | --- | --- |
| **Wheat** | 0.835 ($p < 0.001$) | 0.2158 | -1.3367 |
| **Mouse** | 0.6823 ($p < 0.001$) | -0.9188 | -0.1846 |
| *Arabidopsis* | 0.7547 ($p < 0.001$) | -0.3413 | -0.3758 |
| **Rice** | 0.7808 ($p < 0.001$) | -0.202 | -0.7897 |

**Figure 4.**
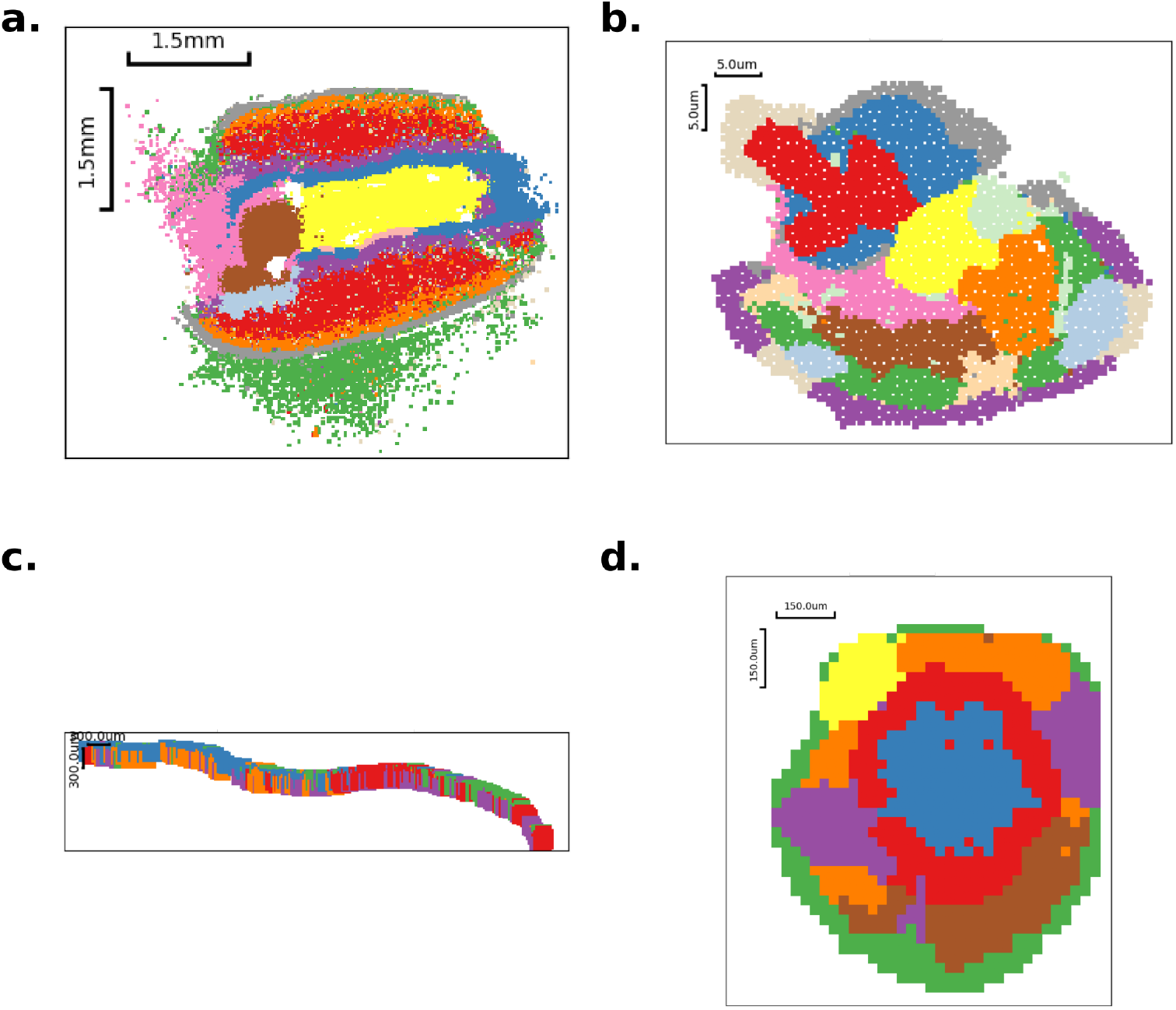
Spatial Leiden clustering across benchmarked spatial transcriptomics datasets. Representative Leiden clustering results generated by the pipeline for the benchmarked datasets: a, wheat; b, mouse; c, Arabidopsis thaliana; and d, rice. These panels illustrate the ability of the pipeline to identify spatially organized transcriptional domains across plant and animal spatial transcriptomics datasets.

The pipeline incorporates a rigorous statistical validation module to assess the distribution of gene expression markers within each identified spatial domain. Across all datasets, the Shapiro-Wilk test yielded values ranging from 0.68 to 0.83 (p<0.05), indicating that the expression profiles follow a non-normal distribution, as expected for spatial transcriptomics data, justifying the use of non-parametric tests in downstream differential expression analysis. Furthermore, the Skewness and Kurtosis metrics provides statistical relevance to the markers, for instance, the wheat dataset exhibited a negative kurtosis (−1.33), suggesting a platykurtic distribution where expression values are spread out. Whereas the mouse dataset showed a higher degree of negative skewness (−0.91), indicating a concentration of highly expressed specific markers. By providing statistical metrics, the pipeline ensures that the user can verify the statistical robustness of the statistically significant clusters.

## Discussion

The pipeline presented here demonstrates the capacity for a robust analysis for ST data. It addresses key challenges related to the data, while also adhering to standardization recommendations required for the development of new technologies, prioritizing utility and reproducibility. A distinguishing feature of the pipeline is its validation on polyploid datasets, providing a benchmarked entry point for plant researchers dealing with complex genomic architectures. By integrating normalization and statistical testing steps, we have made significant progress towards the efficient analysis of complex genomes with ST by preserving and analyzing duplicated gene expression signatures. Furthermore, the pipeline encompasses the generation of co-expression networks, mitigating the analytical challenges of data sparsity and rescuing low-expressed transcripts from being discarded as technical dropouts during network construction [4,35]. By developing sequential steps that take into account the results of previous steps, our pipeline is not only capable of identifying the HVGs and other candidates, but of suggesting the interactions between these and other less expressed genes, commonly removed from spatial analysis, and proposing functional modules and pathways.

A major focus of working with polyploid organisms is dealing effectively with allelic complexity and duplicated genes, a challenge exacerbated by the presence of paralogues. Gene duplications are often drivers of metabolic plasticity, for instance, studies on *Portulaca oleracea* have shown the duplication of *PEPC* genes allow for differential expression to facilitate facultative CAM metabolism[36]. The complexity generated by gene duplications is similarly mirrored in the hexaploid wheat, where the *PEPC* genes consist of multiple isoforms derived from ancient duplications retained across the A, B and D subgenomes [37]. Furthermore, recent spatial transcriptomics analysis of the wheat grain has revealed that these highly similar isoforms exhibit distinct spatial localizations, with specific *PEPC* variants restricted to the pericarp and endosperm, to potentially support C4-like pathways [38]. Accurately profiling and recovering the co-expression networks of highly homologous sequences in spatially resolved data, without losing them to technical dropouts, is critical for interpreting such complex biological mechanisms. Recapitulating these biologically validated spatial distributions demonstrates the reliability and robustness of our pipeline’s sparsity-aware framework.

Current ST research is often fragmented by proprietary technology, which hinders data sharing and meta-analysis [39]. For instance, while 10xVisium provides a framework for exploring tissue architecture, its multi-cell spots require deconvolution to resolve cellular heterogeneity [40,41]. In contrast, Stereo-seq and Slide-seqV2 offer subcellular resolution but generate massive, sparse matrices that demand efficient data handling and aggregation strategies [24,42]. Furthermore, imaging-based methods like MERFISH and Xenium provide subcellular precision, but are often constrained by limited gene panels and lower throughput compared to the unbiased capture of sequencing-based methods [43,44]. Recent studies have introduced sophisticated algorithms targeting specific challenges in high-resolution spatial data, including boundary-assisted single-cell segmentation in plants and animals (STCellbin) [25], adaptive Gaussian smoothing for dropout imputation (EAGS) [11], and graph attention deep subspace networks for spatial domain deciphering (STMSGAL) [27]. However, such methods primarily address isolated steps within the analytical pipeline. Consequently, analyzing ST data still often requires researchers to manually stitch together disparate bioinformatic tools, which frequently leads to significant data loss, user error, and a lack of reproducibility. The primary novelty of the Poly Pipeline lies in overcoming this specific integration bottleneck. Rather than merely acting as a wrapper for existing softwares, it establishes a continuous, precise, and direct workflow. By coupling a universal format converter with multi-mode spatial clustering and sparsity-aware network generation (hdWGCNA), the pipeline minimizes information loss when transitioning between distinct analytical steps. Thus, the pipeline developed addresses a critical gap in spatial transcriptomics by providing a robust, unified solution for the challenges posed by polyploid organisms, mitigating issues of genomic redundancy and data sparsity through advanced normalization and co-expression network analysis.

The pipeline developed addresses a critical gap in spatial transcriptomics by providing a robust workflow validated on complex genomes, minimizing the impact of data sparsity through advanced normalization and co-expression network analysis. Though optimized for polyploid organisms, the pipeline has also proven functional for any organism and, together with its converter, is capable of analyzing any spatial transcriptomics data. By implementing a global pipeline capable of dealing with any type of data input format, our pipeline bridges the gap between different technologies, generating a unified structure promoting the FAIR (Findability, Accessibility, Interoperability, and Reuse) data principles, and serving as a community resource that enhances the reproducibility and utility of spatial transcriptomics [23,45].

## Data availability

The complete pipeline and all the documentation is available in the github page (https://github.com/capuccino26/POLY_PIPELINE) [46]. All the data used in the development and benchmarking are available elsewhere and properly referenced in the text: Wheat:https://www.ncbi.nlm.nih.gov/geo/query/acc.cgi?acc=GSE298021 Rice:https://ftp.cngb.org/pub/stomics/STT0000026/Analysis/STSA0000251/STTS0000395/ Arabidopsis:https://db.cngb.org/stomics/datasets/STDS0000104/ Mouse:https://db.cngb.org/stomics/datasets/STDS0000058/

## Availability of Source Code and Requirements

Project name: POLY_PIPELINE

Project home page: https://github.com/capuccino26/POLY_PIPELINE

Operating system(s): Linux (Optimized for SGE and PBS clusters)

Programming language: Python, R, bash

Other requirements: Python 3.8+, Stereopy 1.5.1, Singularity/Apptainer (for PBS), Conda (for SGE). R libraries: hdWGCNA, WGCNA, Seurat.

License: MIT

RRID: SCR_027993

Bio.tools ID: biotools:poly_pipeline

WorkflowHub DOI: https://doi.org/10.48546/workflowhub.workflow.2091.1

## Authors’ contributions

**Pedro Carvalho:** Writing - Original Draft, Methodology, Software, Validation, Formal analysis, Visualization.

**Tori Millsteed:** Writing - Review & Editing, Conceptualization, Investigation.

**Robert Henry:** Writing - Review & Editing, Conceptualization, Resources, Supervision, Project administration. All the authors read and approved the final manuscript.

**Acknowledgements**

This research was conducted by the Australian Research Council Research Hub for Engineering Plants to Replace Fossil Carbon (project number IH230100006) with funding from the Australian Government.

This study was financed, in part, by the São Paulo Research Foundation (FAPESP), Brasil. Process Number #2025/10055-4.

